# Targeting Thymidine Phosphorylase with Tipiracil Hydrochloride is a Safe and Effective Antithrombotic Therapy

**DOI:** 10.1101/2020.04.25.061234

**Authors:** Abu Hasanat Md Zulfiker, Adam Belcher, Oliver Qiyue Li, Hong Yue, Anirban Sen Gupta, Li Wei

## Abstract

**Rationale:** Most of the current anti-platelet drugs inhibit platelet function permanently and have systemic side effects, including thrombocytopenia and hemorrhage. We previously found that thymidine phosphorylase (TYMP), a platelet cytoplasmic protein, facilitates multiple agonist induced platelet activation and enhances thrombosis. A specific TYMP inhibitor, namely, tipiracil hydrochloride (TPI), has been approved by the U.S. Food and Drug Administration for clinical use as an auxiliary drug making it possible to be repositioned as an anti-platelet medicine.

**Objective:** We aimed to test the hypothesis that TPI is a novel and safe anti-platelet drug by examining its role in platelet activation and thrombosis using both in vitro and in vivo studies.

**Methods and Results:** By co-expression of TYMP and Lyn or Lyn-SH3 domain tagged with glutathione S-transferase, we showed the direct evidence that TYMP binds to the SH3 domain in its partners. TYMP haplodeficiency is sufficient to inhibit thrombosis in vivo regardless of gender. TPI treatment rapidly inhibited collagen- and ADP-induced platelet aggregation, which copied the phenotype of TYMP deficient platelets. Under both normal and hyperlipidemic conditions, treating wild type (WT) mice with TPI via intraperitoneal injection, intravenous injection, or gavage feeding dramatically inhibited thrombosis without inducing significant bleeding. Even administered above the effective dose, TPI has a lower bleeding side effect compared to aspirin and clopidogrel. Most importantly, intravenously delivery of TPI alone or combined with tissue plasminogen activator dramatically inhibited the growth of developing thrombi. Dual administration of very low dose of aspirin and TPI also dramatically inhibited thrombosis without disturbing hemostasis.

**Conclusion:** This pharmacological study demonstrated that TYMP participates in multiple signaling pathways in platelet and plays a mechanistic role in regulating platelet activation and thrombosis. TPI, a specific TYMP inhibitor, would be a novel safe anti-platelet and anti-thrombosis medicine.

## Introduction

Thrombotic events remain a clinically significant area for new mechanistic and therapeutic discoveries, as they are a major cause of morbidity and mortality both in the US and worldwide.^1–3^ However, new anti-thrombotic agents are not being developed since suitable targets are lacking. Platelet activation and hyper-aggregation at the site of vascular injury is the primary pathogenic component of thrombosis, which can lead to vessel occlusion resulting in myocardial infarction and ischemic stroke. In this context, platelet surface receptor glycoprotein VI (GPVI) confers both platelet adhesion and activation in response to exposed collagens at the site of vascular injury.^4–6^ The adhered and activated platelets release soluble agonists, including adenosine diphosphate (ADP), thrombin, and thromboxane A_2_, that activate additional platelets locally via G protein coupled receptors (GPCRs), such as ADP-receptor P2Y12 and thrombin receptor PAR1.^5, 7^ Consequently, various anti-platelet drugs, such as aspirin (COX inhibitor), clopidogrel (P2Y12 inhibitor), and vorapaxar (PAR1 inhibitor), have been developed and used clinically for the primary or secondary prevention of platelet-mediated thrombotic events.^8–11^ However, due to the systemic effects of their targets, these drugs have major side effects including injury to the gastrointestinal mucosa, thrombocytopenia, and systemic hemorrhage.^2, 8–11^ The permanent inhibition on platelet function is also problematic for patients who need emergent surgery.^12^ In addition, some patients do not respond to the current anti-platelet regimens and still have a high incidence of vascular thrombosis.^2^ Therefore, there are significant clinical interests in elucidating unique molecular mechanisms of platelet-mediated thrombus formation, which can lead to the development of a novel anti-platelet therapy with minimized systemic risks.^2, 13^ Currently, all anti-platelet drugs target platelet surface receptors and drugs that safely inhibit platelet function by targeting intracellular proteins have not been developed.

Human thymidine phosphorylase (TYMP) was first isolated from amniochorion^14^ and later from human platelets,^15^ thus it is also known as platelet-derived endothelial cell growth factor.^16^ TYMP is mainly found inside the cell due to the lack of an amino-terminal hydrophobic leader sequence required for cell secretion.^15^ TYMP belongs to the family of pyrimidine nucleoside phosphorylase, and its primary function is to drive the salvage pathway of pyrimidine nucleosides.^17–19^ In the presence of inorganic phosphate, TYMP reversely catalyzes thymidine to thymine and 2-D-deoxyribose-1-phosphate, and the latter is further degraded to 2-D-deoxyribose.^19, 20^ TYMP also has deoxyribosyl transferase activity through transferring the deoxyribosyl moiety from a pyrimidine nucleoside to another pyrimidine base, resulting in the formation of a new pyrimidine nucleoside.^19, 21^ Functional TYMP acts as a homodimer and plays a key role in the pyrimidine nucleoside metabolism, ensuring a sufficient pyrimidine nucleotides pool for DNA repair and replication. However, the role of TYMP in platelets, a non-nucleated cell, remains unclear. Our recent study demonstrated that TYMP is a potential signaling protein and it may transfer cell signaling through the binding of its proline-rich N-terminus to the SH3 domain in its partner proteins.^19, 22^ TYMP deficiency dramatically inhibited platelet response to the conventional agonists, such as collagen, ADP, and thrombin.^22^ Inhibition of TYMP activity with a novel TYMP inhibitor, KIN59,^23^ dramatically inhibited platelet activation in vitro and thrombosis in vivo.^22^ These data suggest that TYMP is a targetable intracellular protein in platelets and its inhibition is antithrombotic. The potent and specific TYMP inhibitor, tipiracil hydrochloride (TPI), is an auxiliary component of a novel anticancer drug, Lonsurf. Lonsurf was recently approved by the FDA for clinical use, suggesting that TPI could be repositioned as a novel antiplatelet and anti-thrombotic drug. In this study, we examined the role of TPI on platelet activation and thrombosis using in vitro and in vivo studies and have provided evidence that TPI-mediated TYMP inhibition is a safe and effective antithrombotic therapy. TPI is suitable as both a primary and secondary prevention for patients with high risk of thrombotic cardiovascular diseases, and combination of low dose TPI and aspirin could be a new dual antiplatelet regimen.

## Methods

The authors declare that all supporting data are available within the article and its online supplementary files.

### Experiment animals

*Tymp*^-/-^ mouse strain was generated by Dr. Hirano’s laboratory as mentioned before.^22, 24^ Wild type C57BL/6 mice (WT) were purchased from Jackson Laboratory (Bar Harbor, ME). All procedures and manipulations of animals have been approved by the Institutional Animal Care and Use Committee of Marshall University (IACUC#: 1033528).

### Murine carotid artery thrombosis model and tail bleeding assay

Detailed ferric chloride (FeCl_3_) induced carotid artery thrombosis model has been described before.^25, 26^ Vessel injury was induced by 7.5% FeCl_3_ solution to the carotid artery for 1 minute. Thrombi formation was observed in real-time using an intravital microscope. The endpoints were set as follows: 1) blood flow has ceased for > 30 seconds; or 2) occlusion is not seen after 30 minutes of FeCl_3_ injury. In this case, 30 minutes was assigned to that mouse for the statistical analysis. In some experiments, a jugular vein catheter prepared with P-10 tubing was placed for drug administration after 5 minutes of thrombus initiation. Thrombus formation was continually monitored using the endpoints mentioned above. Tail bleeding assay was either conducted on mice immediately after the thrombosis study or on mice that were not used for the in vivo thrombosis study.^27^

### Mouse platelet aggregation assay

Mice were anesthetized with ketamine/xylazine (100/10 mg/kg) and whole blood were drawn through inferior vena cava puncture using 0.109 M sodium citrate as an anticoagulant.^22, 28, 29^ Platelet-rich plasma (PRP) was used for platelet aggregation assay. In some experiments, platelets were pretreated with different concentrations of TPI for 2 min (or indicated times) before adding CaCl_2_/MgCl_2_ at a final concentration of 1 mM and agonists as indicated in the Results. Same volume of vehicle (DMSO or saline)-treated platelets was used as controls.

### Cellix flow chamber-based platelet adhesion and aggregation assay

Cellix flow chambers were coated with collagen 10 μg/mL in PBS. Whole blood was drawn from mouse inferior vena cava using 0.109 M sodium citrate as an anticoagulant, stained with Rhodamine 6G, and then used for the flow chamber mediated adhesion and aggregation assay. Some of the whole blood was treated with or without TPI before it was perfused into the chamber.

### Compare the therapeutic effect of TPI to aspirin and clopidogrel

8 to 10 week-old male WT mice were gavage fed with either TPI, aspirin, or clopidogrel once daily for one week, then subsequently used in an in vivo thrombosis study. Tail bleeding assay was conducted on these mice immediately after the thrombosis study.

### Examine the effect of TYMP inhibition on thrombosis under hyperlipidemia

WT mice were fed a western diet (WD, TD.88137) for a total of 4 weeks. For a separate group of mice, the diet was changed to a customized diet (TD.190501), which has the same component as TD.88137 with the addition of 10.7 mg/kg TPI, for the last 7 days of feeding. Mice fed with TD.190501 received approximately 1 mg/kg/day of TPI. These mice were then subjected to the thrombosis study using the 7.5% FeCl_3_ induced thrombosis model.

### Evaluation of platelet signaling activation

Platelets in PRP were stimulated with 2.5 μM ADP for 0, 1, 3, and 5 minutes, then lysed with radioimmunoprecipitation assay buffer containing proteinase/phosphatase inhibitor cocktail. The lysates were subsequently used for AKT activation immunoblotting assays. ADP-stimulated p-selectin expression was analyzed by flowcytometry.

### Generation of fusion proteins to determine that TYMP binds to LYN through the SH3 binding domain

Human SH3 domain nucleotides (hLynSH3, amino acids 63-123) were amplified by PCR and cloned into a mammalian expression vector pEBG (Addgene, plasmid # 22227) with an N-terminal glutathione S-transferase (GST) affinity tag (pEBG-GST-hLynSH3). We had previously constructed a pCDNA6B/his-hTYMP plasmid vector.^30^ pEBG-GST-hLynSH3 and pCDNA6B/his-hTYMP plasmids were co-transfected into COS-7 cells, lysed in a Pierce™ IP Lysis Buffer (ThermoFisher Scientific), and used for GST pull down assay. Eluates were used for western blot assay to confirm the presence of TYMP.

pEGFP-N1-human Lyn-GFP and pCDNA6B/his-hTYMP plasmids were also cotransfected into the COS-7 cells. TYMP was pulled down using immobilized Ni-NTA on magnetic sepharose beads (GE Healthcare Life Sciences) for His-tagged protein purification. His-hTYMP eluates were used for western blot assay for Lyn.

#### Statistics

Data are expressed as mean ± SEM. Results were analyzed by 2-tailed Student’s *t* test or 1-way ANOVA with Bonferroni post-hoc test for multiple comparisons using GraphPad Prism (version 8.3.1). *P< 0.05* was considered statistically significant.

### Supplementary methods

For detailed experimental designs, sample preparation, and experimental conditions, please find the Supplementary Methods provided online.

## Results

### TYMP binds to Lyn SH3 domain

Our previous study demonstrated that TYMP is a novel signaling protein and it binds to the SH3 domain-containing proteins, probably through its proline rich N-terminus.^22^ This finding was confirmed by overexpressing full length human TYMP and Lyn in Cos-7 cells. By pulling down His-Tagged TYMP using the lysates prepared from the co-transfected cells, we found both endogenous and overexpressed Lyn present in the elute (**Fig. 1A**), which corroborates our previous finding using human and mouse platelet lysates. To show the direct evidence, we further generated a GST-SH3 fusion protein and examined its binding to human TYMP by GST pull-down assay. As shown in **Fig. 1B**, pulling down GST-SH3 also pulled down human TYMP. These data showed the direct evidence that TYMP binds to its partner proteins through the SH3 domain.

**Figure 1.**
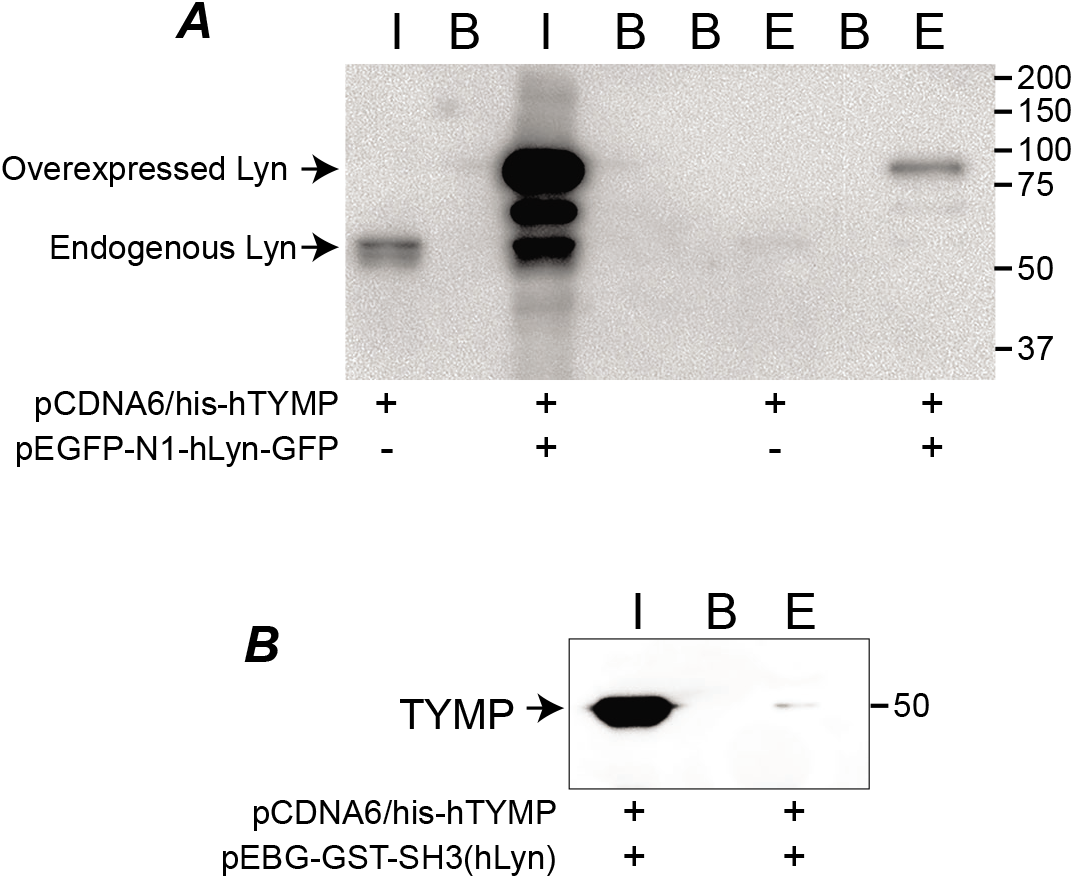
TYMP binds to its partners through their SH3-domain. ***A.*** pCDNA6/his-hTYMP plasmid vector, either alone or combined with pEGFP-N1-hLyn-GFP vector, was transfected into Cos-7 cells, and His-Tagged TYMP was pulled down using His Mag Sepharose Ni beads. Inputs and elutes were blotted using anti-Lyn antibody. ***B.*** pCDNA6/his-hTYMP and pEBG-GST-SH3(hLyn) were co-transfected into Cos-7 cells and the lysate was used for GST pull-down assay and elute was used for blotting human TYMP. In both panels, I: input; B: blank, E: elute. Blots represents 2-3 repeats.

### TYMP-deficiency attenuates platelet aggregation to collagen-coated surfaces

Adhesion to the injury site is an essential function for platelets and is generally viewed as the first step of aggregation, during which specific membrane receptors on the platelet surface binds to cellular and constituents of the extracellular matrix and the vessel wall.^31^ The initial functional receptors primarily include platelet GPIb-XI-V and GPVI. In our previous study, we have found that TYMP deficiency likely does not affect the initial platelet adhesion in the in vivo thrombosis model.^22^ To examine this finding, we coated Cellix Vena8 Fluoro+ chamber with type I collagen,^32, 33^ and perfused the chambers with fluorescently labeled mouse whole blood. As shown in **Fig. 2A**, we found that TYMP-deficiency did not affect platelet binding to the type I collagen-coated surface. However, TYMP-deficiency dramatically decreased platelet aggregation at the end of the three-minute observation. Inhibition of TYMP with 50 μM TPI also significantly reduced platelet aggregation to the collagen-coated surface (**Fig. 2B**). These data suggest that TYMP plays a functional role in GPVI signaling-mediated platelet activation and aggregation; however, TYMP deficiency or inhibition does not affect platelet adhesion to collagen.

**Figure 2.**
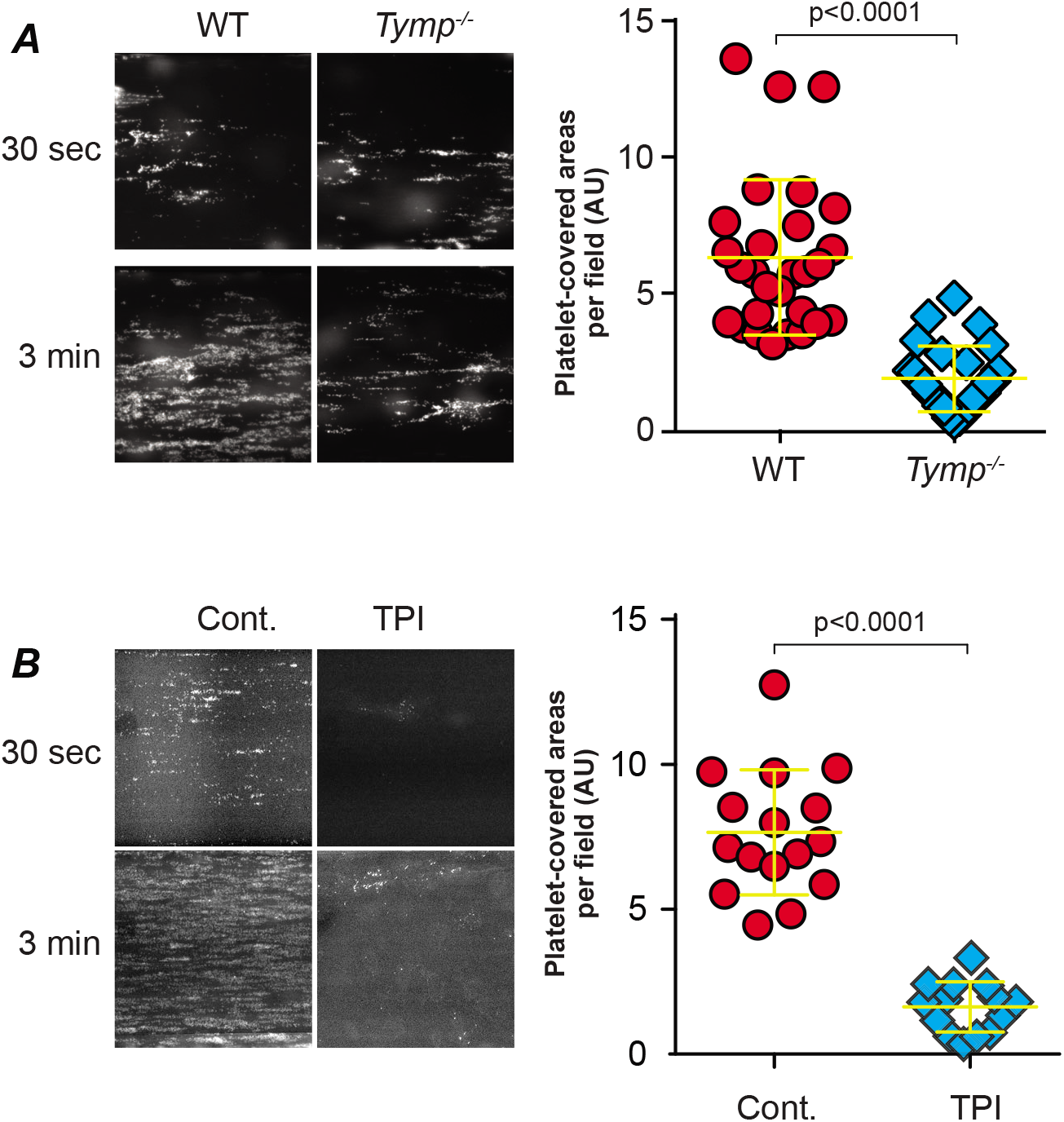
Cellix Vena8 Fluoro^+^ chamber-based platelet aggregation assay. The flow chambers were coated with 10 μg/ml collagen overnight. ***A.*** Whole blood drew from WT and *Tymp*^-/-^ mice were perfused into the chamber in a flow shear 65 dyn/cm^2^. ***B.*** WT whole blood treated with 50 μM TPI in saline or saline alone were perfused into the chamber at a shear 65 dyn/cm^2^. Graphs show areas covered by platelets at 3 minutes after perfusion.

### Tipiracil hydrochloride, a potent and specific TYMP inhibitor, attenuates platelet activation

We have shown that TYMP deficiency significantly attenuated platelet aggregation and P-selectin expression in response to collagen, collagen related peptide (CRP), ADP and thrombin.^22^ TPI, a FDA-approved TYMP inhibitor, recently became commercially available. Therefore, we examined its effects on inhibiting platelet activation and preventing thrombosis. TPI dose-dependently (250, 125, and 62.5 μM) inhibited 0.5 μg/ml CRP-induced platelet shape change (not shown) and 50 μM completely blocked CRP-induced platelet aggregation (**Fig. 3A**). In line with the findings in Fig.2B, pretreatment of platelets with TPI also dose-dependently inhibited collagen-induced platelet aggregation (**Fig. 3B and C**), which phenocopied the behavior of TYMP-deficient platelets. TPI also attenuated ADP-induced WT platelet aggregation but had no effect on *Tymp*^-/-^ platelets (**Fig. 3D**), suggesting that the inhibitory effects of TPI on platelet activation are mediated by TYMP inhibition. This hypothesis is further supported by examination of ADP induced AKT phosphorylation, which has been used as a marker of platelet activation in many studies.^22, 34^ As shown in **Fig. 3E**, we found that ADP-induced phosphorylation of AKT at S473 was bi-phasic. TYMP deficiency likely had no effect on AKT activation within the first minutes after ADP stimulation, but it dramatically delayed the second phase of AKT phosphorylation. In the second phase of AKT phosphorylation, *Tymp*^-/-^ platelets took 10 minutes to reach the AKT phosphorylation levels seen in WT platelets after 5 minutes. Inhibition of TYMP with TPI also dramatically reduced ADP-stimulated AKT phosphorylation, especially the second phase (**Fig. 3F**), and attenuated ADP-stimulated p-selectin expression (**Fig. 3G**).

**Figure 3.**
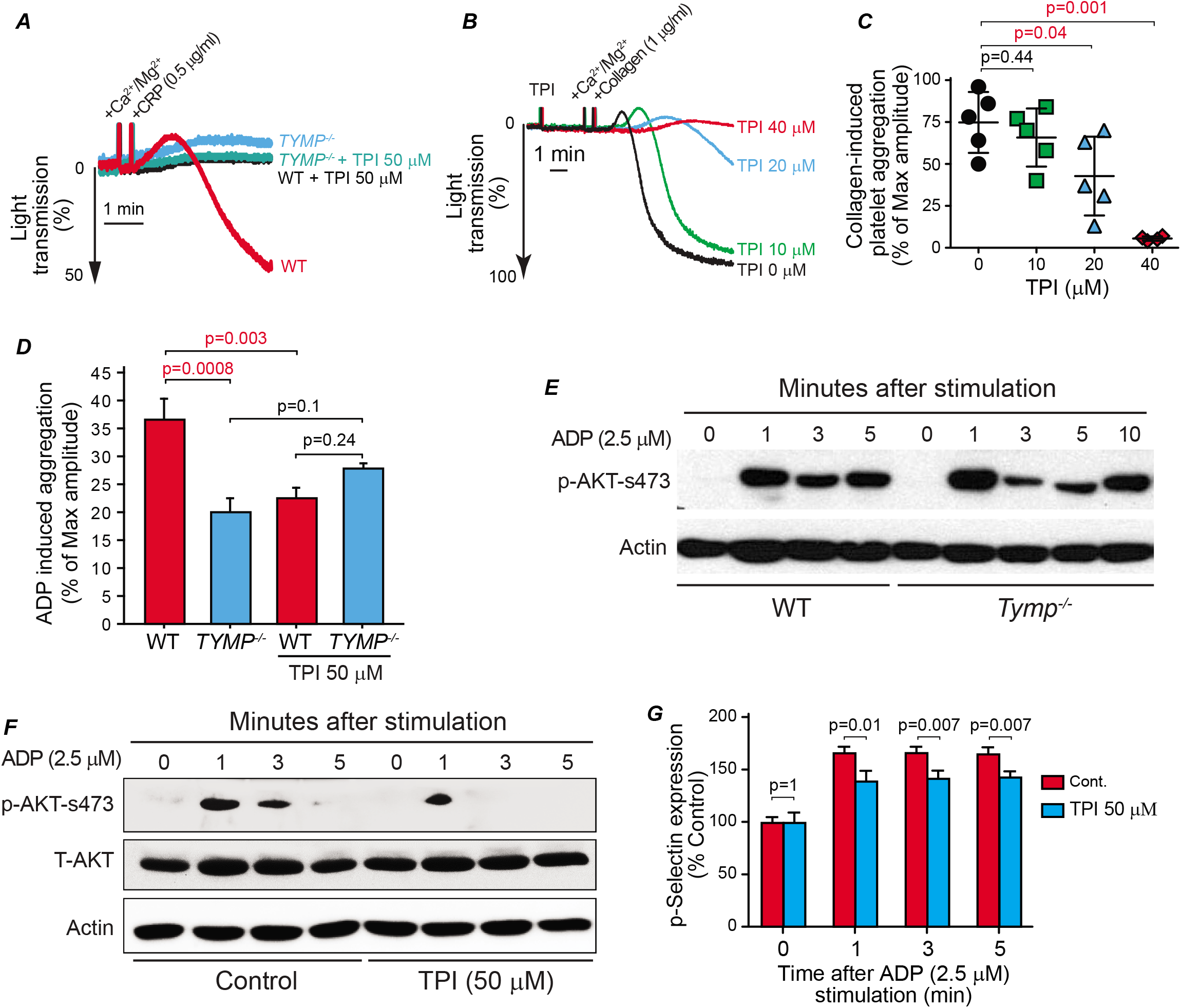
Inhibition of TYMP in vitro inhibits platelet activation. ***A.*** WT and *Tymp*^-/-^ platelets in PRP were treated with 50 μM TPI for 2 minutes and then CRP-induced platelet aggregation was assessed. ***B & C.*** WT platelets in PRP were treated with different concentration of TPI for 2 minutes and then collagen (1 μg/ml) induced platelet activation was assessed. ***D.*** WT and *Tymp*^-/-^ platelets in PRP were treated with 50 μM TPI for 2 minutes and then 2.5 μM ADP-induced platelet aggregation was assessed. ***E.*** WT and *Tymp*^-/-^ platelets in PRP were treated with 2.5 μM ADP for the indicated times and then AKT activation were evaluated. Blot represents two repeats. ***F.*** WT platelets in PRP pooled from 10 mice were divided into 8 parts. Four parts were treated with 50 μM TPI and another 4 parts were treated with saline as controls before they were treated with 2.5 μM ADP for the indicated times. Platelet lysates were used for assessing AKT phosphorylation. ***G.*** WT platelet in PRP were divided into 8 groups, treated with saline or 50 μM TPI, and then treated with 2.5 μM ADP for the indicated times. Platelet surface p-selectin expression was analyzed by flow cytometry and data were shown as ratio of control (0 minute). N=3.

### *Tymp* deficiency results in significant anti-thrombotic effects

We previously reported that TYMP haploinsufficiency or complete deletion significantly inhibited thrombosis in male mice.^22^ We have now found that TYMP deficiency in female mice also significantly inhibits thrombosis (**Fig. 4A and B**). When combining both male and female mice together, we found that cessation of blood flow within 30 minutes was seen in all WT mice (n=17) with an average vessel occlusion time of 9.81 ± 2.25 min (**Fig. 4C and D**). 10 of the 18 *Tymp*^+/-^ mice and 7 of the 16 *Tymp*^-/-^ mice showed flow cessation within the 30 min observation period, with average occlusion times >20 min. These data suggest that TYMP, a platelet cytosolic protein, plays a mechanistic role in platelet activation and thrombosis, which is independent of gender. Partial deficiency of TYMP is enough to achieve a significant antithrombotic effect.

**Figure 4.**
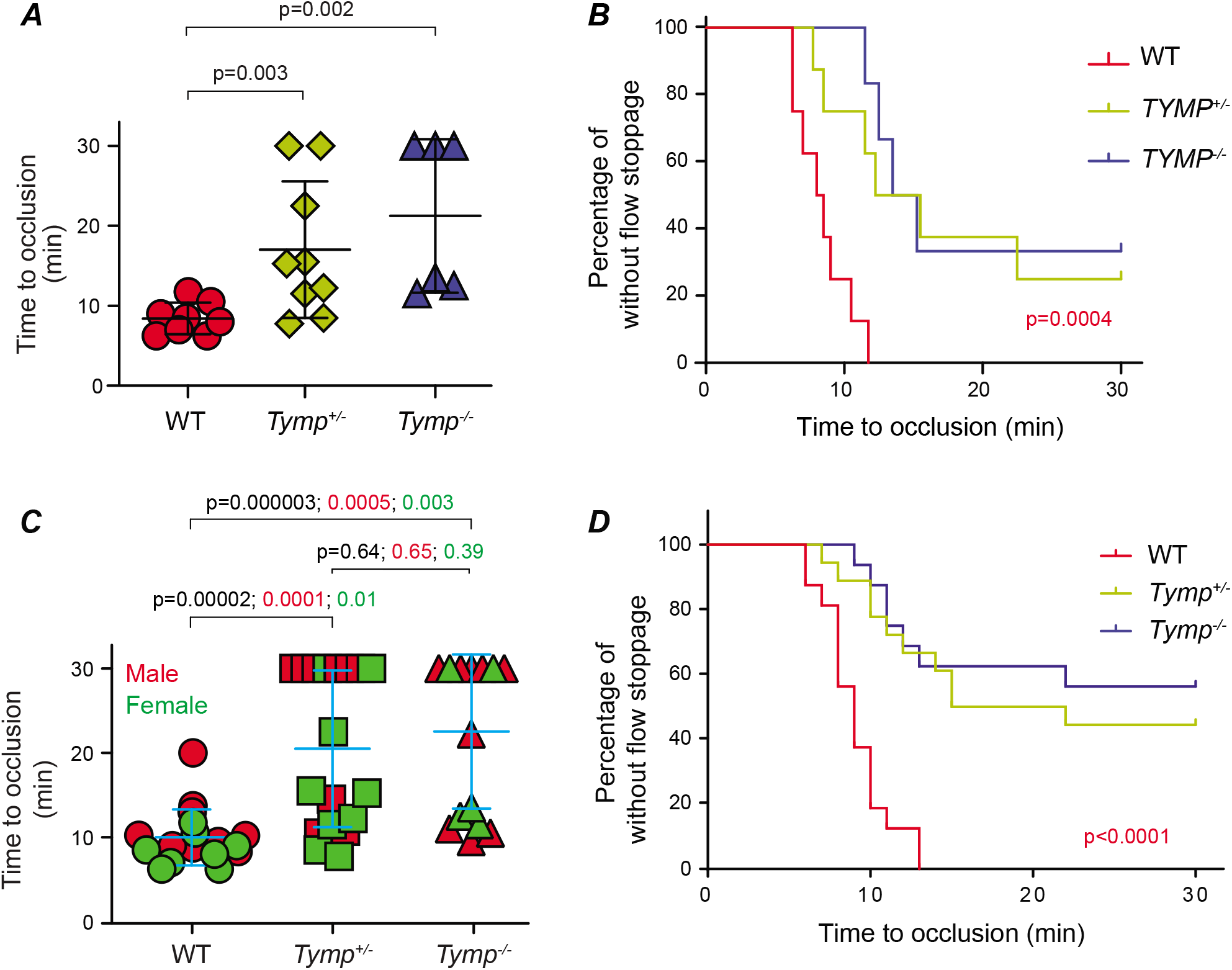
TYMP deficiency in vivo inhibits thrombosis. 8-10 weeks WT and *Tymp*^-/-^ mice in both genders were subjected to the 7.5% FeCl3-induced thrombosis model. ***A & B.*** Role of TYMP deficiency on thrombosis in female mice. ***C & D.*** TYMP deficiency on thrombosis regardless of gender. Time to thrombosis (A and C) and frequency of vessel opening (B and D) were analyzed in WT, *Tymp*^+/-^, and *Tymp*^-/-^ mice.

### TPI inhibits thrombosis under both normal and hyperlipidemia conditions without disturbing hemostasis

Having shown that inhibition of TYMP with TPI significantly inhibits CRP-, collagen-, and ADP-induced platelet aggregation in vitro, we further examined the effects of TPI on in vivo thrombosis. Intraperitoneal injection of TPI once per day for three days at a dose of 17 mg/kg, which, assuming that total body water is 60% of body weight, equals 100 μM plasma concentration, completely inhibited occlusive thrombus formation in the carotid arteries induced by 7.5% FeCl_3_ (**Fig. 5A**). Importantly, this treatment did not significantly affect tail-bleeding time (**Fig. 5B**). Direct intravenous injection of TPI at doses of 1.7 and 0.17 mg/kg, which equals to 10 and 1 μM plasma concentration based on the calculation mentioned above, also significantly inhibited thrombosis (**Fig. 5C**) when compared with mice receiving saline injection alone. Interestingly, gavage feeding of TPI at a lower dose of 60 μg/kg/day, which equals a plasma concentration of 340 nM that is 10-fold higher than the TPI IC50 of 34 nM, for 3 days also significantly inhibited in vivo thrombosis (**Fig. 5D & E**). These data strongly suggest that inhibition of TYMP with its specific inhibitor is an effective antithrombotic therapy.

**Figure 5.**
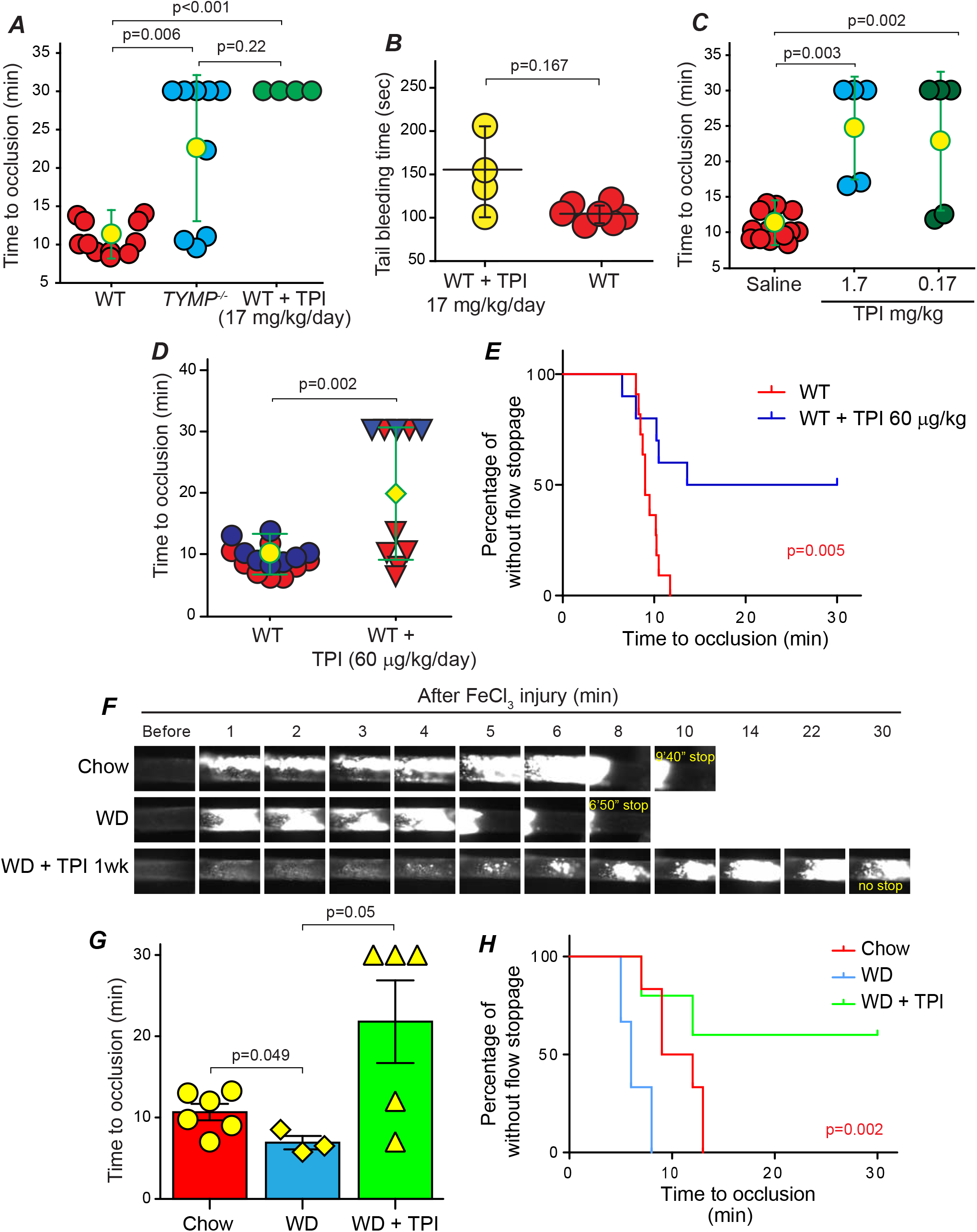
TPI inhibits thrombosis under both normal and hyperlipidemia conditions without disturbing hemostasis. WT mice were treated with TPI by intraperitoneal (IP) injection **(*A*)** or intravenous injection (***C***), and oral administration (***D & E***) at the indicated doses and then subjected to the FeCl_3_ induced thrombosis model. Tail bleeding time was also assessed in mice received TPI IP injection (***B***). ***F, G*** and ***H***, WT mice fed with a western diet (WD, TD.88137) for 4 weeks were subjected to the thrombosis model and thrombosis in age-matched WT mice were used as control. ***F***, representative images from each group; supplementary videos I-III are shown online. ***G***, mean blood flow cessation time. ***H***, Percentage of mice without flow stoppage.

Most of the patients with atherothrombotic vascular diseases have comorbid hyperlipidemia and the associated mortality rates are still unacceptably high. We previously reported that hyperlipidemia leads to platelet hyperactivity, which may contribute to the development of the pro-thrombotic state.^35^ We found that WT mice fed with a WD for 4 weeks is sufficient enough to shorten the thrombosis time when compared to age-matched mice fed with chow. TPI (1 mg/kg) treatment for one week dramatically reduced hyperlipidemia-enhanced thrombosis (**Fig. 5F, G, and H as well online supplementary video I-III**).

### Compare the side effects of TPI vs. aspirin and clopidogrel

Current anti-platelet and anti-thrombotic drugs require chronic dosing to be effective and have bleeding side effects.^2, 13, 36^ Aspirin and clopidogrel are the most frequently used antithrombotic drugs clinically. We thus compared their therapeutic and side effects with TPI. We gavage fed WT mice with different doses of aspirin and clopidogrel for one week, and then subjected them to the 7.5% FeCl_3_-induced thrombosis model. We found that 1 mg/kg/day aspirin and 2.5 mg/kg/day clopidogrel were the lowest effective doses that achieved a significant antithrombotic effect when compared with WT mice without any treatment (**Fig. 6A and supplementary video IV and V**). Although we have found that gavage feeding of TPI, as low as 60 μg/kg/day, significantly inhibited thrombosis in vivo (Fig. 5D & E), we chose a higher dose of TPI, 1 mg/kg/day, in order to explore the potential side effects. We then compared the results to aspirin- and clopidogrel-treated WT mice (**Fig. 6B and supplementary video IV-VI**). When comparing the tail bleeding time, although there were no differences found among the three groups, clopidogrel-treated mice had the longest bleeding time (470 ± 70 seconds), the aspirin-treated mice were second (336 ± 93 seconds), and the TPI-treated mice had the shortest bleeding time (302 ± 67 seconds) (**Fig. 6C**). 75% (3 of the 4) in the clopidogrel group, 50% (3 of the 6) in the aspirin group, and 28.5% (2 of the 7) in the TPI group were considered to have bleeding side effects (**Fig. 6D**). Mice treated with 1 mg/kg/day of TPI had significantly decreased frequency of bleeding than the clopidogrel-treated mice (p=0.034).

**Figure 6.**
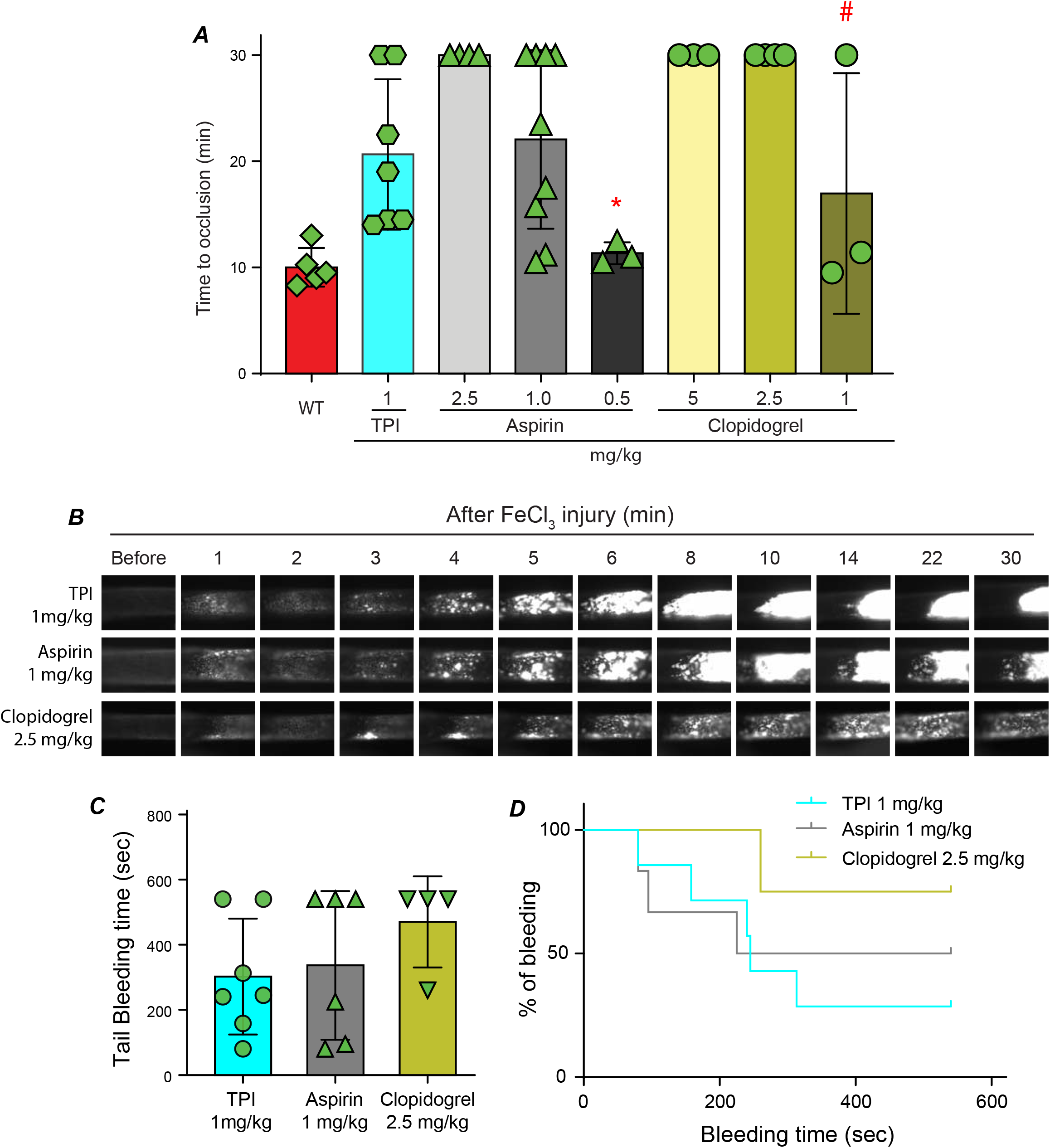
Comparison the therapeutic and side effect of TPI with aspirin and clopidogrel. ***A.*** WT mice were gavage fed with different doses of aspirin and clopidogrel as well as 1 mg/kg TPI in saline and then subjected to the 7.5% FeCl_3_ induced thrombosis model. WT mice received saline were used as control. * & #, p>0.05 vs. WT. ***B.*** Representative video images for mice received 1 mg/kg TPI, 1 mg/kg aspirin, and 2.5 mg.kg clopidogrel. ***C & D*.** Bleeding time in mice received effect doses of aspirin (1.0 mg/kg) and clopidogrel (2.5 mg/kg) was compared with mice received 1 mg/kg TPI.

### TPI is a quick-acting anti-thrombotic drug and co-administration with tissue plasminogen activator (tPA) reduces the tPA dose needed for effective thrombolysis without disturbing hemostasis

The window for treating patients with acute myocardial infarction or ischemic stroke is narrow and tPA is the only FDA-approved clot-busting drug. Intravenous perfusion of tPA has a high risk of systemic coagulopathies and bleeding complications^37^. By using the FeCl_3_-induced carotid artery thrombosis model, we have found that bolus injection of tPA (Activase, 1 mg/kg) 5 min after initiation of vascular injury effectively lysed the established thrombi.^26^ However, due to de novo platelet activation, we also observed persistent formation of new platelet-rich thrombi after tPA administration; thus, thrombi underwent repeated size variations.^26^ By bolus injection of 17 and 1 mg/kg TPI to mice 5 min after 7.5% FeCl_3_-induced vessel injury, we found that thrombosis formation was significantly prolonged or there was no blood flow cessation in those WT mice (**Fig. 7A)**. These data suggest that TPI may be a quick-acting anti-thrombotic drug. We further tested the minimal effective dose of TPI on inhibiting the growth of the on-site thrombus and found that the lowest effective dose is between 50-100 μg/kg. By using the same thrombosis model, we also found that the lowest effective dose of tPA on thrombolysis is between 0.25 to 0.5 mg/kg. Lowering the doses of tPA to 0.25 and 0.125 mg/kg had no thrombolytic action (**Fig. 7B**). Interestingly, when we co-administered TPI 75 μg/kg and tPA 0.25 mg/kg, as shown in **Fig. 7C and 7D**, as well as **supplementary video VII**, we found a significant inhibition of the growth of the on-site thrombi compared to each drug was used individually. Increasing doses of either TPI or tPA in combination also significantly prolonged the thrombosis time (**Fig. 7C**). Most importantly, high doses of TPI (1-4 mg/kg) only slightly prolonged tail-bleeding time to 3.74 ± 0.73 min. These data suggest that co-administration of tPA and TPI reduces the tPA dose required for thrombolysis. TYMP inhibition alone or in combination with a low dose of tPA could be a novel strategy for treating patients with acute myocardial infarction or ischemic stroke without causing coagulopathy.

**Figure 7.**
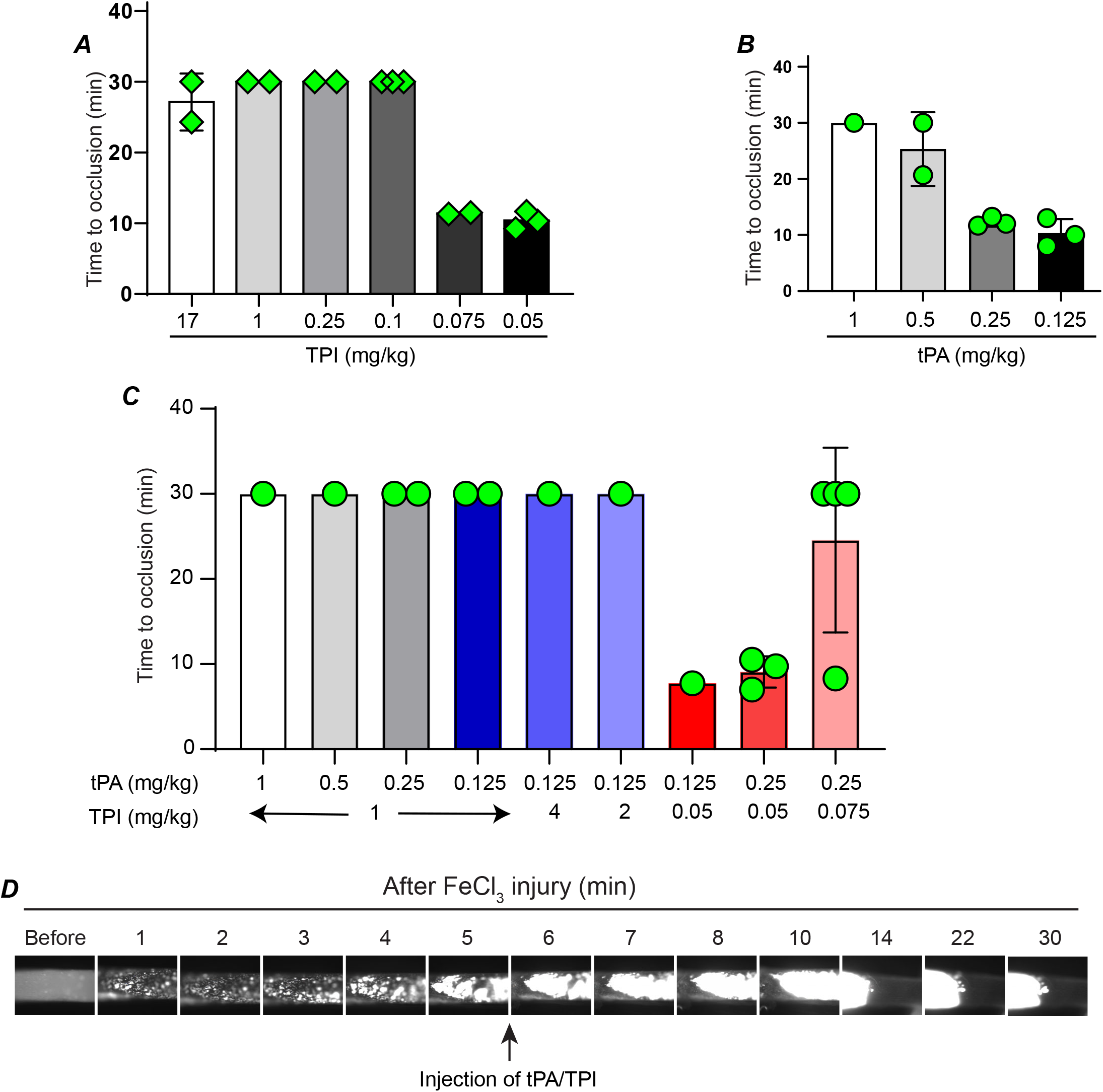
TPI is a quick-acting anti-thrombotic drug that reduces the effective dose of tPA on preventing occlusive thrombi formation. Thrombosis in WT mice were initiated with 7.5% FeCl_3_ and 5 minutes later, TPI (***A***), or tPA (***B***), or the combination of TPI and tPA (***C***) in saline at the indicated doses were bolus injected into mice through a jugular vein catheter, and thrombosis times were assessed. ***D.*** Representative images for mice received 0.25 mg/kg tPA and 75 μg/kg TPI bolus injection.

### Combination of TPI and aspirin is a potential new, safe dual antiplatelet therapy

Dual anti-platelet therapy is commonly used in the first several months after cardiac procedures such as stent placement and transcatheter aortic valve implantation.^38, 39^ Combination of aspirin and P2Y12 inhibitor greatly increases the risk of bleeding, so there is still a need for a robust acute anti-thrombotic treatment with lesser bleeding risk.^40^ We thus examined if combination of aspirin and TPI could be a new dual anti-platelet therapy with neglectable side effect. We chose aspirin 0.5 mg/kg and TPI 50 μg/kg, as treating WT mice with these two drugs alone in the selected dose had no effect on thrombosis (Fig. 6A and 7A). As shown in **Fig. 8A & B**, gavage feeding of WT mice with this regimen significantly prolonged time to occlusive thrombus formation when compared with 0.5 mg/kg aspirin alone. When compared the tail bleeding time with WT mice that were not received any treatment, we found that combination of low doses of aspirin and TPI does not cause bleeding (**Fig. 8C**).

**Figure 8.**
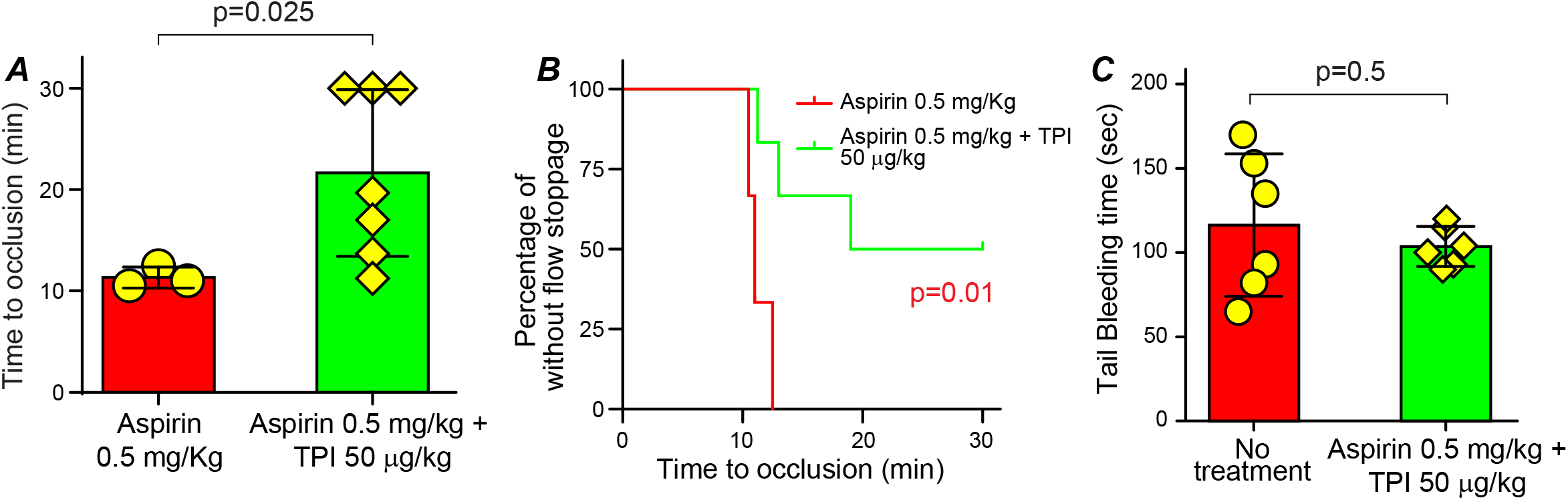
Combination of low dose aspirin and TPI is a new, safe dual antiplatelet therapy. ***A & B*.** WT mice were gavage fed with either aspirin 0.5 mg/kg/day, or a combination of aspirin 0.5 mg-TPI 50 μg/kg/day in 200 μL saline for 7 days. Then the mice were subjected to the 7.5% FeCl_3_ induced carotid artery injury thrombosis model and time to occlusive thrombus formation were determined. ***C*.** Tail bleeding time was assessed using the mice received aspirin 0.5 mg-TPI 50 mg/kg/day after the thrombosis study and compared with age-matched WT mice without any treatment.

## Discussion

Vascular thrombosis is the primary event in life threatening diseases, such as myocardial infarction or ischemic stroke, and platelet activation and aggregation are major components of thrombosis. Consequently, various anti-platelet medications are used clinically for the primary or secondary prevention of thrombosis. However, most of these drugs irreversibly block platelet surface receptors involved in platelet activation and aggregation, which results in systemic side effects, including thrombocytopenia and hemorrhage. Therefore, there is a urgent need to elucidate unique molecular mechanisms of platelet-mediated arterial thrombus formation that can be modulated to allow targeted anti-platelet therapy while minimizing systemic risks^2, 13^. In addition, to our best knowledge, no antiplatelet drugs have been developed to target platelet cytosolic proteins. We recently discovered that TYMP, a platelet cytoplasmic protein, plays important roles in maintaining normal platelet function, and TYMP is also essential for platelet activation induced by multiple agonists, including collagen, ADP, and thrombin.^22^ We have now found that TYMP-enhanced platelet activation is independent of sex and that TYMP deficiency in female mice achieved a similar anti-thrombotic effect to what we saw in male mice.^22^ These new data suggest a universal and mechanistic role of TYMP on platelet activation and thrombosis.

In this study, we thoroughly examined whether TYMP could be inhibited in vivo as well as the consequences following TYMP inhibition using its specific and potent inhibitor, TPI. TPI is water soluble (>13.25 mg/ml). These is no published study examined the pharmacokinetics of TPI alone. Administration of a single dose of Lonsurf at 35 mg/m^2^ generates the absorption rate of TPI of area under curve is 301 ngoh/ml. The maximum plasma concentration of TPI is 69 ng/ml, which was observed at 3 h after administration.^41^ At steady state, TPI half-life is 2.4 h, and around 77% is excreted through faeces and urinary with unchanged structure.^42^ TPI was delivered to mice using three different routes, including intraperitoneal injection, intravenous injection, and oral administration, and the effect was tested using different disease models. Our data clearly demonstrated that TPI is an effective and safe anti-thrombotic compound, even under a hyperlipidemic condition. The effective dose of TPI could be as low as 60 μg/kg/day without significantly affecting hemostasis. Most importantly, intravenous injections of TPI, either delivered alone or in combination with tPA, also significantly inhibited the growth of the ongoing thrombi, which makes TYMP inhibition a potential alternative remedy for patients with acute myocardial infarction or stroke. This fast-acting characteristic also indicates that TPI can be used for percutaneous coronary intervention and transcatheter aortic valve implantation etc, either alone or combined with other drugs to temporally inhibit platelet function, as TYMP inhibitors rapidly and reversibly inhibit platelet aggregation.^22^

TYMP has a pro-angiogenic effect and previous studies of TYMP are primarily focused on its function on endothelial cells^16, 43–45^ and cancer angiogenesis.^46–48^ Platelets are a major source of TYMP, with each human platelet containing about 5,400 to 11,600 copies of TYMP,^49, 50^ but the function of TYMP in platelet physiology and function remains unclear. TYMP has been implicated in diseases that have high risk of thrombosis, such as atherosclerosis,^51, 52^ cancer,^53, 54^ and diabetes mellitus.^55^ In addition, ionizing radiation, which induces significant TYMP expression, has been associated with thrombotic vascular occlusion.^56^ In the current study, we found that TYMP deficiency or inhibition does not affect platelet binding to the collagen-coated surface; however, it dramatically attenuated platelet aggregation on the collagen-coated surface in a flow chamber assay. In combination with our published data,^22^ these data further demonstrated that TYMP likely plays a more important role in the GPVI signaling pathway. GPVI is found exclusively on platelets and megakaryocytes and is the predominant platelet receptor for collagen.^57^ Deficiency of GPVI in humans and in mice is not associated with a strong bleeding diathesis.^58^ Hitherto, there is no effective and convenient inhibitor targets GPVI. A recent phase I study using Revacept, a dimeric GPVI-Fc fusion protein that blocks collagen/GPVI binding mediated platelet activation, demonstrated that targeting the GPVI signaling is a safe intervention.^59^ However, Revacept is delivered by intravenous injection, therefore has limited situational use, such as percutaneous coronary intervention.^60^ Revacept has no effects on circulating resting platelets, which indicates that it may have no effect on hyperactive platelets under certain diseased conditions. In contrast, our data suggest that TPI may have more broad indications than the other anti-platelet drugs.

In addition to GPVI signaling, TYMP also participates in GPCR-mediated platelet activation. In our previous study, we showed that TYMP deficiency reduced ADP-induced platelet p-selectin expression. Here, we further demonstrated that TYMP deficiency or inhibition attenuated ADP-induced AKT phosphorylation in platelets. ADP-induced platelet activation requires concomitant signaling from both P2Y1 and P2Y12 receptors that couple to Gαq and Gαi, respectively. However, ADP-induced AKT activation is predominantly mediated by the platelet P2Y12 receptor.^61^ While we still do not know how TYMP affects GPCR signaling pathways, TYMP deficiency may put a “brake” on autocrine or paracrine mechanisms of platelet GPCR activation.

These data provide a strong rationale that TPI could be repositioned as a potential antithrombotic medicine. To this end, we compared the prophylactic therapeutic effects and side effects of TPI with aspirin and clopidogrel using the murine FeCl_3_ induced thrombosis model. We found that the lowest dose for aspirin and clopidogrel in inhibiting mouse thrombosis is 1 mg/kg and 2.5 mg/kg, respectively. As predicted, these doses of aspirin and clopidogrel dramatically prolonged bleeding time. Although a high dose of TPI (1mg/kg) occasionally prolonged bleeding time in some mice, mice that received doses of TPI (60 μg/kg) showed no prolonged bleeding time in all cases (**Supplementary Figure I**). As an auxiliary component of the anti-cancer drug, Lonsurf (TAS-102), TPI has been evaluated in clinical trials ^62–65^ and has been shown to be systemically safe. Neither systemic bleeding nor mitochondrial neurogastrointestinal encephalomyopathy, an extremely rare autosomal recessive disease that is reportedly associated with *TYMP* gene loss-of-function mutations, was observed. TYMP deficiency does not affect high dose thrombin-stimulated platelet activation.^22^ These data suggest that TPI may be safer and more beneficial than the current anti-platelet drugs. Most importantly, oral administration of very low doses of aspirin and TPI dramatically inhibited thrombosis without inducing any coagulopathy, suggesting that this combination could be a novel remedy for patients with high risk of thrombosis. Clinical trial studies are necessary to demonstrate this hypothesis.

In summary, our study demonstrated that TYMP, a platelet cytosolic protein, plays an important role in platelet activation and thrombosis. TYMP can be safely and rapidly inhibited by TPI. TPI-mediated TYMP inhibition dramatically inhibited platelet activation and thrombosis in both normal and hyperlipidemic conditions, as well as in different disease models. Under effective anti-thrombotic doses, TPI does not cause bleeding disorders, as found in patients treated with other anti-platelet drugs.

## Supporting information

Supplemental video I-chow

Supplemental video II-western diet

Supplemental video III-western diet + TPI

Supplemental video IV-aspirin

Supplemental video V-clopidogrel

Supplemental video VI-TPI

Supplemental video VII-tPA+TPI

## Non-standard Abbreviations and Acronyms

GPVI: glycoprotein VI
ADP: adenosine diphosphate
GPCRs: G protein coupled receptors
TYMP: thymidine phosphorylase
TPI: tipiracil hydrochloride
WT: wild type
FeCl_3_: ferric chloride
PRP: platelet-rich plasma
GST: glutathione S-transferase
CRP: collagen related peptide
tPA: tissue plasminogen activator

## Acknowledgements

No other persons besides the authors have made substantial contributions to this manuscript.

## Sources of Funding

This study is supported by Marshall University Institute Fund (To Dr. Wei Li), Marshall University, School of Medicine and College of Pharmacy Collaborative Grant (PI: Dr. Wei Li), NIH R15HL145573 (PI: Wei Li), NIH R01HL129179 (PI: Anirban Sen Gupta, Co-I: Wei Li), NIH R01HL130090 (PI: Thomas M McIntyre, Co-I Wei Li) and WV-INBRE grant P20GM103434 (PI: Gary Rankin).

## Disclosures

None

## List of supplemental materials

Expanded Materials & Methods

Online Video 1 – VII

Online Figure I

Legend for supplemental figure and videos.

## SUPPLEMENTAL MATERIAL

### Materials

All platelet agonists including ADP, collagen and thrombin was purchased from Chronolog (Havertown, PA). Collagen related peptide (CRP) was a gift from Dr. Peter Newman (Blood Research Institute, WI). Antibodies to phosphorylated AKT and total AKT were purchased from Cell Signaling Technology (Dallas, TX). FITC-conjugated P-selectin antibody and the isotype control IgG were purchased from **BD Biosciences** (San Jose, CA). All other chemical reagents were purchased from Sigma (St. Louis, MO) except where specifically indicated.

### Experiment animals

Wild type C57BL/6 mice (WT) were purchased from Jackson Laboratory (Bar Harbor, ME). *Tymp*^-/-^ mouse strain was generated by Dr. Hirano’s laboratory as mentioned before,^1,2^ and maintained in our lab by inbreeding. Breeder mice were genotyped before mating and all mice were given identity using ear tags at the time of weaning. Mice in eight to twelve weeks old in both genders were used in this study. All procedures and manipulations of animals have been approved by the Institutional Animal Care and Use Committee of Marshall University (IACUC#: 1033528).

### Murine carotid artery thrombosis model and tail bleeding assay

Detailed ferric chloride (FeCl_3_) induced carotid artery thrombosis model has been described before.^3,4^ Briefly, the mouse was anesthetized with ketamine (100 mg/kg) and xylazine (10 mg/kg). Vessel injury was induced by topically applying a piece of filter paper (1 x 2 mm) saturated with 7.5% FeCl_3_ solution to the carotid artery for 1 minute. Thrombi formation was observed in real-time using an intravital microscope (Leica DM6 FS) and video images were captured with a 14-bit Retiga R1 CCD color digital camera (Teledyne QImaging, Surrey, BC, Canda) and a STP7-S-STDT Streampix 7 software (Norpix, Montreal, Canada). The endpoints were set as: 1) blood flow has ceased for > 30 seconds; or 2) occlusion is not seen after 30 minutes of FeCl_3_ injury. In this case, 30 minutes was assigned to that mouse for the statistical analysis.

In some experiments, a jugular vein catheter prepared with P-10 tubing was placed for drug administration after thrombus initiation. In this case, platelets were labeled by perfusion of 50 μL Rhodamine 6G solution through the jugular vein catheter and then thrombosis was initiated as mentioned above. TPI (62.5 - 2,000 μg/kg), tPA (125-1,000 μg/kg), or combination of different doses of TPI and tPA were delivered to the mice 5 minutes after thrombus initiation. Thrombus formation was continually monitored using endpoints as mentioned above.

Tail bleeding assay was conducted in some mice immediately after the thrombosis study as well as in some mice that were not used for the in vivo thrombosis study.^5^ Briefly, the tail was cut at the site of 1 cm from the tip with a sharp razor blade and immediately immersed into warm saline (37 °C) and cessation time of bleeding was recorded. If bleeding showed no stoppage after 9 minutes, which is about three times longer than the average bleeding time that we observed in the normal WT mice, 9 minutes was assigned to that mouse for statistical analysis. Red blood cells collected from the bleeding assay were separated by centrifugation, lysed in RBC lysis buffer, and hemoglobin amount was measured with a wavelength of 550 nm.

### Mouse platelet isolation

Mice were anesthetized with ketamine/xylazine (100/10 mg/kg), and whole blood were drawn through inferior vena cava (IVC) puncture using 0.109 M sodium citrate as an anticoagulant.^1,6^ Modified Tyrode’s buffer (in mM: 137 NaCl, 2.7 KCl, 12 NaHCO3, 0.4 NaH2PO4, 5 HEPES, 0.1% glucose and 0.35% BSA, pH 7.2), 0.7 volumes of the whole blood, was added, and platelet-rich plasma (PRP) was separated by centrifugation at 100 *g* for 10 min, and transferred into a new tube. The buffer coat and red blood cells sedation were further centrifuged at 1,000 g for 6 minutes to obtain platelet-poor-plasma (PPP). Platelets number were counted with a hemocytometer.

### Platelet aggregation assay

Murine platelets in PRP were adjusted to a concentration of 2.5 × 10^8^/ml using PPP. Platelet aggregation was performed using a standard turbidimetric assays monitored by an Aggregometer (Model 700, Chrono-log, Havertown, PA) and data were acquisted using Aggrolink8 version 1.29.^6–8^ Sample were stirred constantly at 1,200 RPM at 37 °C. Light transmission was monitored over time, and aggregation was quantified, with 100% aggregation corresponding to 100% light transmission. In some experiments, platelets were pretreated with different concentration of TPI for 2 min (or indicated times) before adding CaCl_2_/MgCl_2_ for a final concentration of 1 mM and agonists as indicated in the Results. Same volume of vehicle (DMSO or saline)-treated platelets was used as controls.

### Cellix flow chamber-based platelet adhesion and aggregation assay

Cellix flow chambers were coated with collagen 10 μg/mL in PBS for overnight and then blocked with 0.1% BSA in PBS for 30 minutes at room temperature. Whole blood was drawn from mouse IVC using 0.109 M sodium citrate as an anticoagulant and stained with Rhodamine 6G in a final concentration of 0.1 μg/μL. The whole blood was repleted with CaCl_2_/MgCl_2_ in a final concentration of 1 mM immediately before perfused through the flow chamber. In some experiments, whole blood drew from one mouse was divided into two parts (450 μL/part), and one part was treated with TPI in a final concentration of 50 μM for 5 minutes, repleted with CaCl_2_/MgCl_2_, and then perfused into the chamber. Another part treated with saline was used as control. The flow shear was set at 65 dyn/cm^2^. Platelet accumulation on the coated chamber surface was monitored and recorded in real time for 3 minutes with an Invert Microscope (Leica DM8) with a video camera operated by the CaptaVision+ v2.1 software (ACCU-SCOPE®, Commack, NY). The flow chambers were washed with PBS for additional 3 minutes to remove the free blood cells and then images at the six marker-sites designed by the manufacture were taken and fluorescent intensity was measured using Image J (Fiji, NIH).

### Compare the therapeutic effect of TPI with Aspirin and clopidogrel

8 to 10 week old male WT mice was gavage fed with TPI (0.06, 0.17, and 1 mg/Kg), Aspirin (0.5, 1, and 2.5 mg/Kg), or clopidogrel (1, 2.5, and 5 mg/Kg), or the combination of aspirin 0.5 mg/kg and TPI 50 μg/kg, once daily for one week, and then the mice were subjected to the thrombosis model. Tail bleeding assay was conducted on these mice immediately after the thrombosis study. In another set of study, WT mice received intraperitoneal injection of TPI 14 mg/kg or vehicle for 3 days, or gavage feeding with TPI (0.06 mg/Kg/day) for 14 to 30 days, and then the mice were subjected to the thrombosis model and tail bleeding assay.

### Examine the effect of TYMP inhibition on thrombosis under hyperlipemia

Increased platelet reactivity and thrombosis in dyslipidemia has been observed in multiple murine models of in vivo thrombosis as well as in clinical patients. To determine whether TPI attenuates the hyperlipidemia-enhanced thrombotic diathesis, we fed WT mice with a western diet (TD.88137) for 4 weeks to enhance hyperlipidemia. For some mice, the diet was changed to a customized diet TD.190501, which has the same component as TD.88137 but contains TPI 10.7 mg/Kg, for the last 7 days. Mice fed with TD.190501 received TPI in a dose of about 1 mg/Kg/day. These mice were then subjected to the thrombosis study using the 7.5% FeCl_3_ induced thrombosis model. Age-matched mice on chow diet were used as control.

### Immunoblotting assay-based evaluation of platelet signaling activation

Platelets in PRP were pooled from 8-10 mice, and platelet concentration was adjusted to 2.5 × 10^8^/ml with PPP and then divided into 4 equal parts. CaCl_2_/MgCl_2_ in a final concentration of 1 mM was added immediately before platelets activation was initiated with 2.5 μM ADP. Platelet activation was stopped by adding an EDTA/PGE1 (2 mM and 1μg/ml final concentration, respectively) mixture at 1, 3, and 5 min. No ADP-treated platelets were used as resting platelet control. Platelets were pelleted immediately by centrifugation and lysed in radioimmunoprecipitation assay (RIPA) buffer (ThermoFisher Scientific, Waltham, MA) containing proteinase/phosphatase inhibitor cocktail (ThermoFisher Scientific). Thirty micrograms of total proteins were separated in SDS-PAGE, transferred to a PVDF membrane, and blotted with antibodies as indicated in the Result. Membranes were stripped and re-blotted with an actin antibody as loading control.

### Flowcytometry assay of platelet activation

Platelets in PRP were adjusted to 2.5 × 10^8^/ml with PPP and then divided into 8 equal parts. CaCl_2_/MgCl_2_ in a final concentration of 1 mM was added immediately before platelets activation was initiated with 2.5 μM ADP. Platelet activation was stopped by adding an EDTA/PGE1 (2 mM and 1μg/ml final concentration, respectively) mixture at 1, 3, and 5 min. No ADP-treated platelets were used as resting platelet control. Platelets were then fixed with 2% paraformaldehyde, and then stained with a FITC conjugated rat anti-mouse p-selectin antibody (BD pharmingen) and analyzed by flowcytometry.

### Generation of fusion proteins to determine that TYMP binds to LYN through the SH3 binding domain

We previously showed that TYMP binds to Lyn, Fyn and Yes, and this binding may be via the proline-rich N-terminus of TYMP to the SH3 domain in the SFKs.^1^ To further test this hypothesis, the human SH3 domain nucleotides (hLynSH3, amino acids 63-123) was amplified by PCR using primers 5’-ATCGTGGTCGCCCTGTAC-3’ and 5’-ATTCAGCTTAGCCACGTAGTTAGAT-3’ and a codon optimized human Lyn encoding plasmid pEGFP-N1-human lyn-GFP (Addgene, plasmid # 35958) as a template. The hLynSH3 PCR product was cloned into a pCR2.1 TA cloning vector (Invitrogen, Carlsbad CA), transformed into One Shot TOP10 *E. coli* (Invitrogen), and screened with blue and white selection. The insert orientation was confirmed by PCR sequencing with the M13 forward primer, 5’-GTAAAACGACGGCCAGT-3’, and a reverse primer 5’-ATCGTGGTCGCCCTGTAC-3’. The hLynSH3 domain was further cloned into a mammalian expression vector pEBG (Addgene, plasmid # 22227) with an N-terminal glutathione S-transferase (GST) affinity tag between the restriction sites BamHI and NotI, and a new plasmid pEBG-GST-hLynSH3 was generated.

We had previously constructed a pCDNA6B/his-hTYMP plasmid vector.^9^ pEBG-GST-hLynSH3 and pCDNA6B/his-hTYMP plasmids were co-transfected into COS-7 cells using X-tremeGENE 9 DNA transfection reagent (Roche, Mannheim Germany). The cells were harvested 36 hours later and lysed in a Pierce™ IP Lysis Buffer (ThermoFisher Scientific). One milligram of total protein was added to Glutathione MagBeads (GenScript, Piscataway, NJ) and purified by affinity chromatography based on the manufacture’s introduction. The GST-hLynSH3 eluates was used for western blot assay to confirm the presence of TYMP.

pEGFP-N1-human Lyn-GFP and pCDNA6B/his-hTYMP plasmids were also cotransfected to the COS-7 cells. TYMP was pulled down using immobilized Ni^+^ on magnetic sepharose beads (GE Healthcare Life Sciences) for His-tagged protein purification. To do this, the cell lysates were adjusted to contain (in mM) 5 imidazole, 200 Na3PO4, and 500 NaCl, then adjusted to a pH of 7.4 before added to the His Mag Sepharose Ni beads. His-hTYMP eluates were used for western blot assay for Lyn.

#### Statistics

Data were expressed as mean ± SEM. Results were analyzed by 2-tailed Student’s *t* test or 1-way ANOVA with Bonferroni post-hoc test for multiple comparisons using Graphpad Prism (version 7.03). *P< 0.05* was considered statistically significant.

---

**Supplemental figure I.**
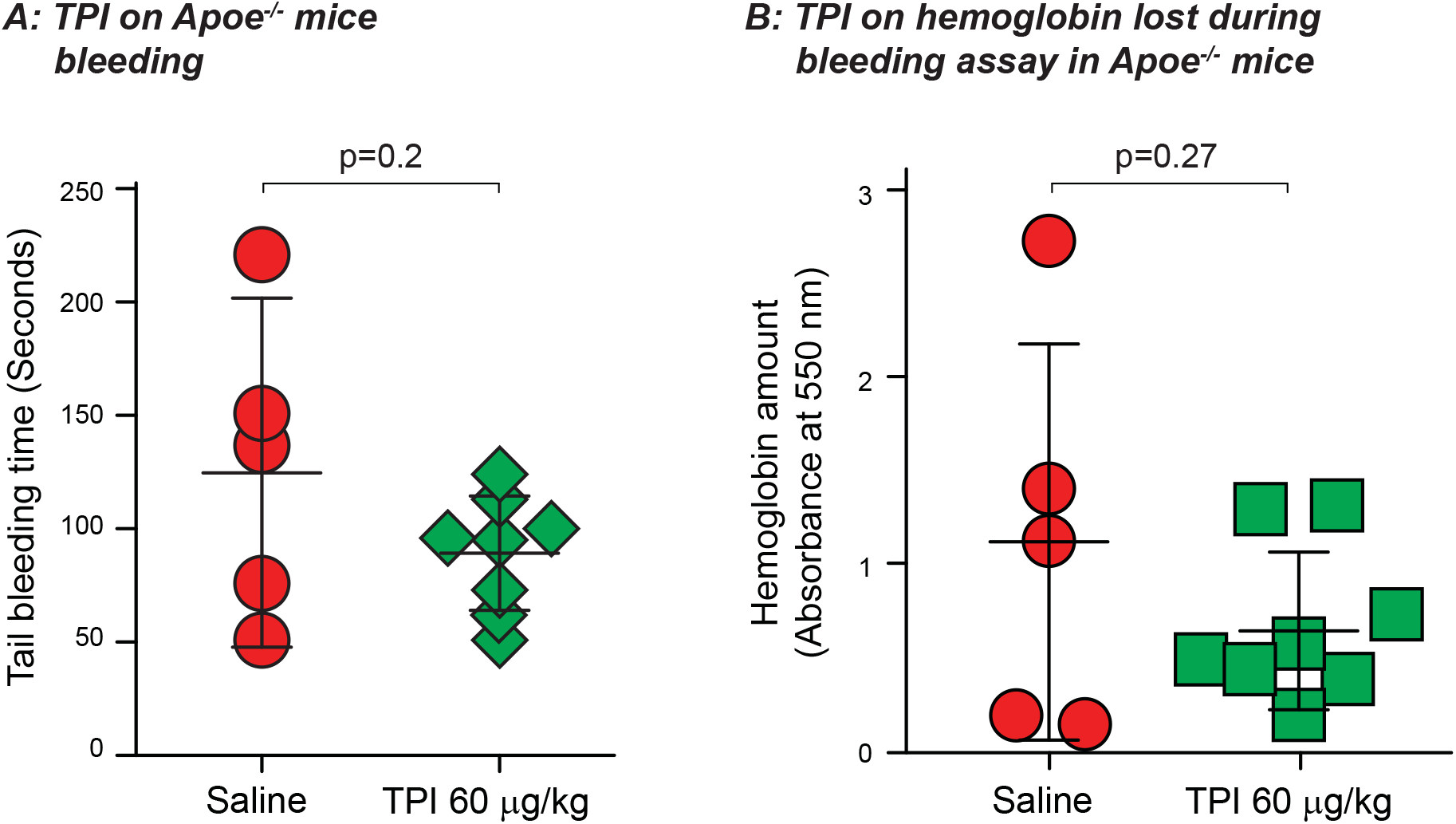
*Apoe*^-/-^ mice in 8 weeks old were fed with a western diet (TD.88137) and at same time, the mice were gavage fed with TPI at a dose of 60 mg/kg for 5 weeks. ***A*.** Tail bleeding time was conducted by cutting 1 cm length of tail and immersed immediately in 37 °C saline. ***B*.** Red blood cells were collected by centrifuge, lysed in RBC lysis buffer, and hemoglobin amount was evaluated by measuring wavelength absorbance at 550 nm.

## Legends for online supplementary videos

**Supplemental video I:** Thrombosis formation in female mice fed with chow.

**Supplemental video II:** Thrombosis formation in female mice fed with a western diet (TD.88137) for 4 weeks.

**Supplemental video III:** Thrombosis formation in female mice fed with a western diet (TD.88137) for a total of 4 weeks. From the 4^th^ week, mice received a customized diet TD.190501, which has the same component as TD.88137 with the addition of 10.7 mg/kg TPI. Mice fed with TD.190501 received approximately 1 mg/kg/day of TPI.

**Supplemental video IV:** Thrombosis formation in male WT mice fed with 1 mg/kg aspirin for 1 week.

**Supplemental video V:** Thrombosis formation in male WT mice fed with 2.5 mg/kg clopidogrel for 1 week.

**Supplemental video VI:** Thrombosis formation in male WT mice fed with 1 mg/kg TPI for 1 week.

**Supplemental video VII:** Thrombosis formation was initiated in male WT mouse with 7.5% FeCl_3_ treatment for 1 minute. TPI 75 μg/kg and tPA 0.25 mg/kg in 100 μL saline was perfused to the mouse through a jugular vein catheter 5 minutes after vessel injury. Thrombus formation was observed for a total of 30 minutes.

## Notes

### Competing Interest Statement

The authors have declared no competing interest.

